# Scaling Functional Annotation Across Proteomes, Pangenomes and Metagenomes with Sma3s v3

**DOI:** 10.64898/2026.09.14.748778

**Authors:** Alejandro Rubio, Jesus L. García-Junco Alcalá, Elisa Luque-Jiménez, Angel Martin Dominguez, Joaquín Dopazo, Antonio J. Perez-Pulido, Carlos S. Casimiro-Soriguer

## Abstract

High-throughput sequencing has generated protein datasets whose scale increasingly exceeds the practical limits of conventional functional annotation workflows. We present Sma3s v3, a scalable reimplementation of the Sma3s three-step annotation strategy, which combines transfer from highly similar homologs, orthology-based inference, and functional enrichment among homologous proteins. Sma3s v3 replaces BLAST-based searches with MMseqs2 and introduces parallel processing, reusable SQLite caches, taxonomic filtering, and traceable outputs that retain the evidence underlying each assignment.

We evaluated the method on a Vibrio cholerae pangenome comprising 50,415 gene clusters from 11,295 quality-filtered genomes and on a metagenomic catalogue containing 843,935 proteins. After excluding non-informative assignments, Sma3s v3 annotated 30,662 pangenome clusters (60.8%), comparable to InterProScan (60.2%) and exceeding eggNOG-mapper (41.4%), while providing 5,747 annotations not recovered by either comparator. Gene Ontology comparisons showed broad semantic agreement between methods, with Sma3s v3 frequently contributing more non-redundant information in Molecular Function and Biological Process. Within the pangenome, annotation coverage reached 97.1% for core clusters and approximately 59% for accessory and unique clusters. Exact protein matches to non-Vibrio genera identified 1,838 candidate horizontally transferred clusters enriched in genetic mobility, antimicrobial resistance, and metal tolerance functions.

In the metagenomic catalogue, Sma3s v3 annotated 728,014 proteins (86.3%), compared with 616,895 (73.1%) using InterProScan 2026, and recovered approximately 20,000 unique functional terms. These results establish Sma3s v3 as a scalable and interpretable tool for functional annotation and re-annotation of proteomes, pangenomes, and metagenomic protein catalogues.

## Introduction

The advent of high-throughput sequencing has transformed molecular biology and computational microbiology, making the generation of genomes, transcriptomes, and metagenomes routine. However, this increase in scale has not been accompanied by an equivalent capacity to experimentally characterize the function of proteins encoded by these data. Automatic functional annotation has therefore become an essential step in transforming sequences into biological knowledge, as it allows genes and proteins to be linked to molecular functions, biological processes, cellular components, and metabolic pathways^1,2^. The significance of this process stems from the fact that a substantial portion of subsequent analysis is contingent upon the quality, coverage, specificity, and consistency of the initial annotations.

The Sma3s method was originally developed as a three-step modular system for the automated functional annotation of large sets of proteins and protein-coding sequences, particularly complete proteomes and transcriptomes. This system utilizes a simple tool with low computational requirements. Previous versions of the software enabled users to assign gene names, protein descriptions, Gene Ontology (GO) terms, EC numbers, pathways, and UniProt keywords, as well as to generate functional summaries that were useful for comparing organisms^3,4^. The enduring relevance of the second version of Sma3s is evident not only in its documented use but also in its sustained citation impact. According to Web of Science (accessed August 2026), the publications describing the first and second versions have received 53 and 81 citations, respectively. Recent studies have employed Sma3s in various biological contexts, including bacterial genome and pangenome annotation^5,6^, plant metabolic reconstruction and developmental transcriptomics^7,8^, single-cell transcriptomics of helminths^9^, stress response transcriptomics in dinoflagellates^10^, and a cross-domain analysis of 1,020 proteomes encompassing 12.8 million proteins^11^. Nevertheless, the contemporary state of functional genomics has undergone a substantial transformation.

In recent years, the unit of analysis has expanded beyond the genome, proteome, or transcriptome of a single organism. In the contemporary realm of comparative genomics, there has been a marked shift towards the utilization of pangenomes, metagenomic catalogs, genomes assembled from metagenomes, and extensive collections of proteins derived from intricate microbial communities^12,13^. The concept of the pangenome facilitates the representation of the complete repertoire of genes in a population or clade, thereby enabling the distinction between the core genome, which comprises genes conserved in all or most of its members, and the accessory genome, which consists of genes present only in a subset of them. Core genes have been observed to exhibit greater levels of conservation, resulting in their more extensive representation in databases. In contrast, accessory genes have been shown to demonstrate a higher degree of variability. These accessory genes have been implicated in ecological adaptations, phenotypic diversity, genetic mobility, and horizontal gene transfer^14^.

Horizontal gene transfer is a primary driving force in the evolution of prokaryotic genomes and is closely linked to the dynamics of the accessory genome. Large-scale comparative studies have demonstrated that the genes implicated in horizontal transfer are predominantly accessory^15^. This underscores the necessity for tools capable of efficiently annotating not only conserved and easily recognizable genes, but also variable, rare, or potentially mobile genes that may help explain adaptive differences among strains, species, or communities. In this context, functional annotation facilitates the interpretation of the pangenome’s composition and the characterization of genes previously identified as candidates for horizontal gene transfer, including those with potential ecological, clinical, or biotechnological relevance.

In the context of metagenomics, catalogs derived from microbial communities encompass a multitude of genes and proteins from both cultured and uncultured organisms. These catalogues are frequently characterized by incomplete assemblies, fragmented sequences, or genomes assembled from metagenomes. For instance, recent catalogs of the gut microbiome during the early stages of life have compiled tens of thousands of metagenome-assembled genomes (MAGs) and more than 80 million protein sequences^16^. Analogously, the investigation of environmental metagenomes has led to the identification of hundreds of thousands of novel gene families, predominantly associated with uncultivable taxa. A significant proportion of these gene families have yet to be functionally characterized, yet they manifest evolutionary and ecological signals indicative of their relevance^17^. The functional annotation of these repertoires, therefore, necessitates methods capable of processing large volumes of sequences and retrieving useful information even when there is limited similarity to previously characterized proteins.

The current functional annotation ecosystem comprises a variety of powerful tools, although they were designed with different objectives and trade-offs. InterProScan functions as a centralized resource for the classification of proteins into families, domains, and functional sites by integrating multiple signature databases, with mappings to GO terms, structures, and other resources^1^. EggNOG-mapper v2 has demonstrated the efficacy of orthology-based functional transfer as a strategy for large genomes and metagenomes^18^. In the domain of bacterial genome annotation, Bakta integrates the identification of genomic elements with conventional functional annotation methods^19^. Other tools, such as DRAM and HUMAnN, address more specific needs related, respectively, to microbial metabolism reconstruction and functional community profiling^20,21^. This diversity is indicative of methodological fragmentation, wherein certain tools prioritize the depth and diversity of annotations, while others prioritize speed or scalability. Additionally, there are tools that prioritize metabolic reconstruction, comprehensive genomic annotation, or community profiling. Consequently, a single strategy is not universally optimal for all data and biological questions.

This fragmentation is further compounded by a semantic challenge. The function of protein is not adequately represented by a single label; rather, it constitutes a structured, hierarchical description whose degree of specificity can vary considerably. Gen<u>e</u> Ontology has been identified as the most widely used framework for annotating sequences, and it continues to be updated regularly with new revisions of the ontological graph^2^. Consequently, the evaluation of a functional annotation tool should not be confined to the percentage of annotated proteins or the exact match of terms and it must also consider the specificity, consistency, and semantic similarity of the annotations. This notion is consistent with the principles established by CAFA, which has underscored the necessity to rigorously evaluate functional prediction methods. Furthermore, it is in alignment with the recent advancements in deep learning-based models and protein language models, such as DeepGO-SE, which integrate sequence, ontological knowledge, and neuro-symbolic learning to enhance functional prediction^22,23^.

For many experimental researchers, functional annotation is not only a matter of assigning ontology terms, but also of obtaining informative gene names, protein descriptions, and complementary functional information that enables rapid biological interpretation. Since its first release, Sma3s has been designed with this objective in mind, providing rich, interpretable annotations that facilitate the rapid characterization of genes and proteins in newly generated datasets. In this paper, we present Sma3s v3, an evolution of Sma3s designed to address this new scenario. Following the initial release of its first versions, Sma3s has gained widespread adoption within the scientific community. This sustained adoption underscores the efficacy of its original approach, which is predicated on ease of use, minimal computational requirements, and the generation of interpretable functional results. However, it also highlights the necessity to adapt the tool to the prevailing increase in the scale and complexity of omics data. In this regard, Sma3s has been modified to enable scalable functional annotation of extensive protein sets in comparative genomics, pangenomics, and metagenomics contexts. Sma3s v3 incorporates an updated architecture and more efficient search methods, with the objective of improving performance on massive datasets without compromising functional annotations useful for subsequent analyses. In addition, the performance of the tool was evaluated in comparison to established benchmark tools. Its utility in biologically challenging scenarios was also explored, including the annotation of pangenomes, accessory genes, horizontal gene transfer candidates, and metagenomic data. Consequently, Sma3s v3 signifies a technical update to a well-established tool and its adaptation to the shift in scale experienced by functional genomics. The scope of research encompasses proteome and transcriptome annotation, as well as the interpretation of gene repertoires derived from complex microbial communities.

## Materials and Methods

### Implementation of Sma3s v3

The implementation was developed in Python 3.11.15 using only modules from the standard library, including sqlite3 for storing intermediate data on disk. Sequence similarity searches were performed using MMseqs2 v18.8cc5c via easy-search command^24^, which replaced the BLAST-based searches utilized in previous versions.

The UniProtKB bacterial and archaeal files corresponding to version 2026_02 were utilized as reference databases in the analyses of this work^25^. When UniProt files were provided in “.dat” format, Sma3s generated the corresponding FASTA and “.annot” files, extracting protein sequences and functional fields, including gene name, protein description, EC numbers, Gene Ontology terms, UniProt keywords, and metabolic pathways. To optimize this parsing, parallelizable architecture was implemented, limited to one-quarter of the maximum number of threads to prevent RAM saturation. Optionally, parameters were included to select or exclude reference entries based on UniProt’s taxonomic hierarchy at the genus, family, or order level. In addition, Sma3s generated reusable SQLite cache for FASTA offset indexes, processed annotations, MMseqs2 hits, and best reciprocal hit maps to reduce memory usage and avoid reprocessing large files.

To assess the efficacy of the strategy, the initial MMseqs2 search was executed with a maximum E-value of 1e-6, a sensitivity of 7.5, low-complexity masking disabled by default, and a maximum of 250 hits retained per sequence. The resulting hits were then evaluated sequentially by the three Sma3s annotators: A1, based on highly similar homologs; A2, based on orthologous sequences; and A3, based on functional-term enrichment among homologous proteins. For Annotator 1, a minimum identity of 90%, a minimum reference sequence coverage of 90%, and a p-value ≤0.1 were required. For Annotator 2, the following criteria were met: a best reciprocal hit ratio, a minimum reference sequence coverage of 80%, and a p-value ≤0.1. The identity threshold was calculated using the Rost curve with a parameter of 20. Annotator 2’s reciprocal search was redesigned to avoid executing separate searches for each hit. Consequently, it performs batch reciprocal searches using MMseqs2 stored in SQLite indexes, which are accessible during the final annotation transfer phase. For Annotator 3, instances with identity higher than the Rost curve were considered, and the final annotations were selected using a hypergeometric test with a p-value ≤0.1. The homologous sequences from Annotator 3 were clustered using the MMseqs2 easy-cluster method with the following parameters: 95% identity, 95% coverage, and --cov-mode 0. These thresholds have been incorporated as configurable parameters.

Subsequent to the search with MMseqs2, the final annotation for each protein was parallelized using the concurrent.futures.ProcessPoolExecutor function. Each worker established independent connections to the SQLite cache for hits, annotations, the FASTA index, and best reciprocal hits. They then applied the logic of the three annotators, prioritizing Annotators 1 and 2 over Annotator 3, to the assigned proteins. Finally, they returned annotation records and partial counts to the main process. The results were merged by constructing a table containing each protein’s original identifier and the transferred functional fields.

The Sma3s v3 and its associated auxiliary scripts were disseminated through a Conda recipe. The recipe encompassed the primary annotation program, ancillary scripts for output processing and functional analysis, executable entry points, and the dependencies necessary for their execution, including Python 3.11 and MMseqs2. The package under consideration was built using the conda-build tool, with the specific versions of the dependencies and the files installed in the environment taken into account. The installation was configured to be performed from the PMC_fps channel using the command "conda install -c PMC_fps -c conda-forge -c bioconda Sma3s", so that the main program and auxiliary utilities would be available from the command line in a controlled environment^26^.

### Creation of a Pangenome for *Vibrio cholerae*

The complete set of assembled *Vibrio cholerae* genomes available in the National Center for Biotechnology Information’s Genome Database as of January 2026 was collected, including complete and draft genomes^27^. The total number of genomes downloaded was 16,618. Duplicate genomes and those with a mean Average nucleotide identity (ANI) below 90% relative to the rest of the dataset, calculated using skani version v0.3.2^28^, were removed. Subsequently, assemblies exhibiting more than 1% gaps or with NG50 values below the first quartile calculated from the subset of lower-quality genomes were discarded. These values were calculated using QUAST v5.3.0 tool^29^.

The prediction of protein-coding genes was conducted using Bak<u>t</u>a v1.12^19^. The pangenome was constructed via Panaroo v1.7.0, with an identity threshold of 90% and the --merge_paralogs parameter to prevent the separation of paralogs at that threshold^30^. Additionally, Panaroo was executed with the following supplementary options: The parameters in question are len_dif_percent, clean-mode sensitive, and remove-invalid-genes. Clusters containing sequences recovered during Panaroo’s correction process were removed from the final matrices.

### Annotation and functional enrichment of the *Vibrio cholerae* pangenome

Protein clusters in *V. cholerae* pangenome were classified according to their frequency of occurrence in the final set of analyzed genomes. Clusters that were present in ≥95% of the genomes were defined as core; those present in a single genome were defined as unique; and clusters present in more than one genome but in <95% of the set were defined as accessory. The representative protein sequence for each cluster was annotated using Sma3s v3 with the UniProtKB bacterial database as a reference. A cluster was designated as annotated if it exhibited at least one functional field retrieved by Sma3s, encompassing gene name, protein description, EC number, Gene Ontology terms, UniProt keywords, or functional pathways.

Concurrently, representative sequences from the pangenome were annotated using eggNOG-mapper v2.1.15, with MMseqs2 serving as the search engine^18^. The analysis was constrained to the bacterial domain, and a maximum e-value of 1×10-5 was employed, along with a minimum coverage of 50% for both the query and the target. Subsequently, the proteins were analyzed with InterProScan v5.78-109.0, employing all available databases to obtain annotations for domains, families, and protein signatures, as well as GO terms and associated functional pathways using the -goterms and -pa options^1^.

The assessment of functional enrichment was conducted by employing the auxiliary script designated as "functional_enrichment.py", which is incorporated within the Sma3s v3 conda recipe. This assessment was performed in a distinct manner for both UniProt keywords and Gene Ontology terms. GO terms were subsequently categorized into three distinct categories: biological process, molecular function, and cellular component, through the implementation of the GO-basic.obo database. For each pangenome category, each term was compared against the rest of the pangenome using a one-sided exact Fisher’s test. The p-values were corrected using the Benjamini–Hochberg method. The concept of enrichment was operationalized through the calculation of fold enrichment, which is defined as the proportion of clusters annotated with a term within a category divided by the corresponding proportion in the entire pangenome. Terms that received support from fewer than five clusters were excluded from further consideration. In the unique group, overly general GO terms were removed only from the graphical panels using a curated exclusion list, while the complete, unfiltered tables were retained as the analysis output.

### Functional and Semantic Comparison of Annotation Tools

The annotations generated by Sma3s, eggNOG-mapper, and InterProScan were integrated using pangenome cluster identifiers as common identifiers. For each tool, the sources of functional evidence available in their output files were extracted. The technical fields that were derived from the search or alignment were not regarded as direct functional evidence. Prior to the comparison, non-informative annotations were filtered out. A comprehensive review of the existing literature revealed the exclusion of several categories of proteins from the analysis. These categories include "single description", hypothetical proteins, uncharacterized proteins, unnamed products, terms with unknown functions, DUF/UPF domains, and the COG S category.

In Sma3s, the following were considered: gene name, protein description, EC number, GO terms, UniProt keywords, and functional pathways. In eggNOG-mapper, the following were considered: preferred name, functional description, GO terms, EC numbers, KEGG pathways and reactions, COG categories, CAZy, KEGG-TC, and BiGG reactions. In InterProScan, the following elements were taken into consideration: member databases, accessions and signature descriptions, InterPro accessions, GO terms, and functional pathways.

The evaluation of functional coverage was conducted on a cluster level, with the recording of the presence or absence of at least one valid source of functional evidence for each tool. The comparison was performed independently for its three main ontologies using Gene Ontology (GO) terms. For each gene, tool, and ontology, the assigned GO terms were retrieved and collapsed to reduce hierarchical redundancy. To achieve this objective, ancestral terms were eliminated when a more specific descendant term was present within the same set. The remaining terms were grouped into semantic clusters using Wang’s method, a measure based on the structure of the GO graph that quantifies the similarity between two terms based on overlap and the weighted semantic contribution of their ancestral terms^31^. Terms exhibiting a Wang similarity score of at least 0.90 were designated as belonging to the same semantic cluster. Subsequently, a single representative term was selected from each cluster for further analysis.

The semantic alignment between tools was calculated for each gene and ontology by comparing the collapsed sets of representative GO terms using the Best Match Average (BMA) strategy^32^. For each term in one set, the term in the other set that exhibited the highest Wang similarity was identified, and the BMA score was obtained by averaging these best matches in both directions. The resulting Wang/BMA score, ranging from 0 to 1, reflected the degree of semantic agreement between annotators after reducing the internal hierarchical redundancy of their GO annotations. In independent, in pairwise comparisons, the annotator with the greater number of representative semantic clusters was considered to provide greater effective semantic diversity. In instances where both annotators identified the same number of clusters, the case was designated as equivalent in terms of information content, though not necessarily in terms of semantic content. Genes exhibiting a Wang/BMA similarity score below 0.90 were designated as cases of low semantic agreement between tools.

Tables were processed using the following packages: readr v2.2.0, dplyr v1.2.1, tidyr v1.3.2, stringr v1.6.0, purrr v1.2.2, and tibble v3.3.1^33^. The GO ontology structure was obtained using GO.db v3.23.1, and semantic similarity was calculated using GOSemSim v2.38.3^34^. The figures were generated using ggplot2 v4.0.3, and the intersection diagrams were generated using ComplexUpset v1.3.3^35^.

### Identification and functional enrichment of candidate genes for horizontal gene transfer (HGT)

Candidate genes for horizontal gene transfer were defined as clusters in the *Vibrio cholerae* pangenome whose representative sequence showed 100% amino acid identity and 100% coverage when compared to proteins from genera other than Vibrio, based on the annotation obtained using Sma3s v3. These clusters were integrated with the pangenome classification to determine their distribution among the core, accessory, and unique categories. The non-*Vibrio* genera associated with perfect matches were summarized by the number of candidate clusters. These clusters were interpreted as associated taxa rather than definitive donors. The functional enrichment of the candidate genes was evaluated using the "functional_enrichment.py" auxiliary script, with the complete pangenome serving as the background and applying the same statistical framework described for pangenome functional enrichment.

### Functional annotation and analysis of metagenomic proteins

In order to evaluate Sma3s v3 in a metagenomic context, the public MGYA00131824 analysis from MGnify (SRA: SRR4101290), associated with study MGYS00001842, was utilized^36^. The available assembled protein sequences, including both previously annotated and unannotated proteins, were downloaded and analyzed collectively as a single metagenomic protein catalogue. The present dataset was annotated using the Sma3s v3 and Interproscan v5.78-109.0 tools, with the same parameters described in the previous section on functional annotation.

The functional recovery was evaluated using analytical rarefaction of unique functional terms. For each annotator, the terms considered were restricted to the same information sources represented in the global annotation panel. For a term present in *m* proteins within a universe of *N* proteins, the expected probability of observing it when sampling n proteins was calculated as *1 - C(N - m, n) / C(N, n)*. The expected number of unique functional terms for each sample size was obtained by summing this probability across all terms detected by each method.

The taxonomic signal associated with metagenomic proteins was estimated based on the best Sma3s hits. Hits with ≥90% amino acid identity and ≥90% coverage of the query protein were designated as high-confidence hits. The identifiers of the best hits were then mapped against the taxonomy associated with the reference proteins, and the number of proteins assigned to each species was counted. These counts were utilized as a high-confidence protein similarity signal, rather than as a direct estimate of taxonomic abundance.

## Results

### Sma3s v3 retains the original annotation logic and incorporates a scalable architecture

The latest version of Sma3s has been developed as a modular reimplementation, with the objective of enhancing the speed, scalability, and reproducibility of the protein functional annotation process. The tool maintains the foundational philosophy of Sma3s v2, which employs a progressive annotation strategy utilizing three modules or annotators: annotation transfer from highly similar homologs (A1), inference from orthologous sequences (A2), and functional assignment based on terms enriched in groups of homologous sequences (A3). This approach facilitates a balance between annotation coverage and reliability, a central principle of the original Sma3s design (Figure 1A).

**Figure 1.**
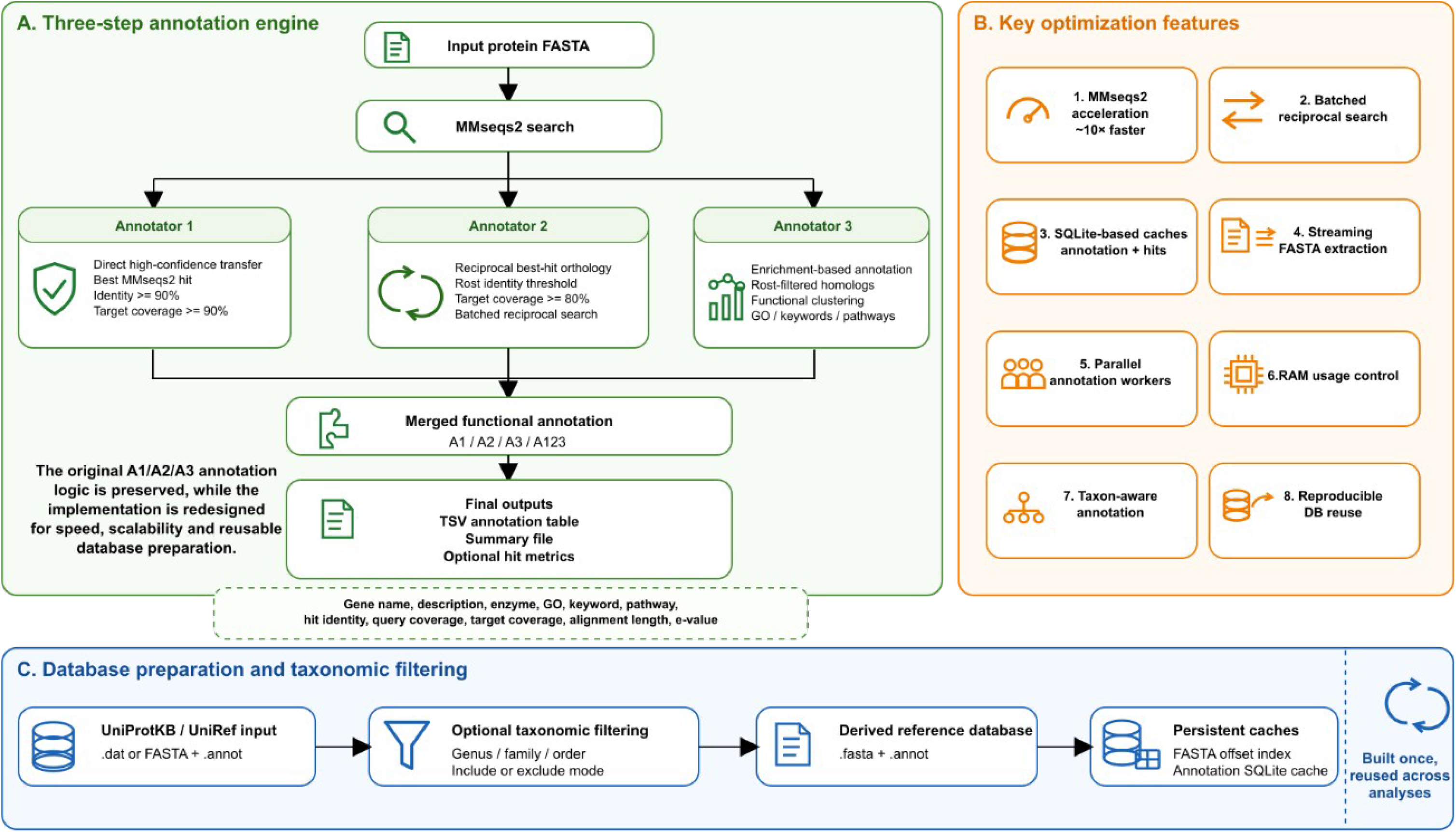
Architecture and optimizations of Sma3s v3. A) Schematic representation of the three-step annotation engine, which remains unchanged from Sma3s v2 but has been substantially accelerated. The input protein sequences are then compared against the reference database using MMseqs2 and processed by three complementary annotators: direct transfer from high-confidence hits, assignment based on best reciprocal hits, and consensus/enrichment annotation from multiple homologs. The annotations generated by the three modules are integrated into a final functional annotation, including gene name, description, EC number, GO terms, keywords, functional pathways, and alignment metrics when requested. B) Major computational improvements incorporated into Sma3s v3 to increase the performance and stability of the annotation workflow. C) Reference database preparation workflow. Sma3s is a software that can be used to process various data files. It can accept UniProtKB/UniRef files in ".dat" format or FASTA databases accompanied by their ".annot" file. It also allows for the application of optional taxonomic filters by genus, family, or order. Furthermore, Sma3s generates derived databases along with persistent caches of FASTA indexes and SQLite annotations. These resources are constructed on a single occasion and subsequently reused in subsequent analyses.

In order to enhance scalability, Sma3S v3 incorporates an SQLite-based intermediate storage strategy for the creation of reusable caches (Figure 1C). This architecture serves to minimize the requirement to load substantial structures into memory, while concomitantly circumventing the need for repetitive parsing and indexing of the same databases in successive iterations. Furthermore, parallelization has been integrated at several pivotal points, including the parsing of input files and the final annotation process (Figure 1B).

Additionally, Sma3s v3 enhances the traceability of the annotation generated. The tabular output maintains the conventional functional categories of Sma3s, encompassing gene name, description, EC, GO, keywords, and metabolic pathways. Moreover, it can incorporate the annotator responsible for each assignment and the hit metrics employed to transfer the annotation. This information enables users to differentiate between annotations derived from direct hits, reciprocal relationships, or consensus among homologs, thereby facilitating the subsequent application of filters based on identity, coverage, or annotation source. In essence, Sma3s v3 upholds the foundational principles of Sma3s while offering an enhanced architecture that is both more efficient and reproducible in its functional annotation of proteomes, pangenomes, and assembled metagenomic data.

To facilitate the installation and reproducible use of Sma3s v3, the tool was packaged along with its auxiliary scripts into a Conda recipe. This distribution enables the concurrent installation of both the primary annotation program and the ancillary utilities for result processing and functional analysis within a unified environment. After the implementation of the PMC_fps channel, the Sma3 executables become accessible directly from the command line. This development eliminates the necessity for manual installation of dependencies and concomitantly reduces variability across systems. This strategy facilitates the execution of the complete annotation workflow and derived analyses on workstations or high-performance computing environments (Figure 2).

**Figure 2.**
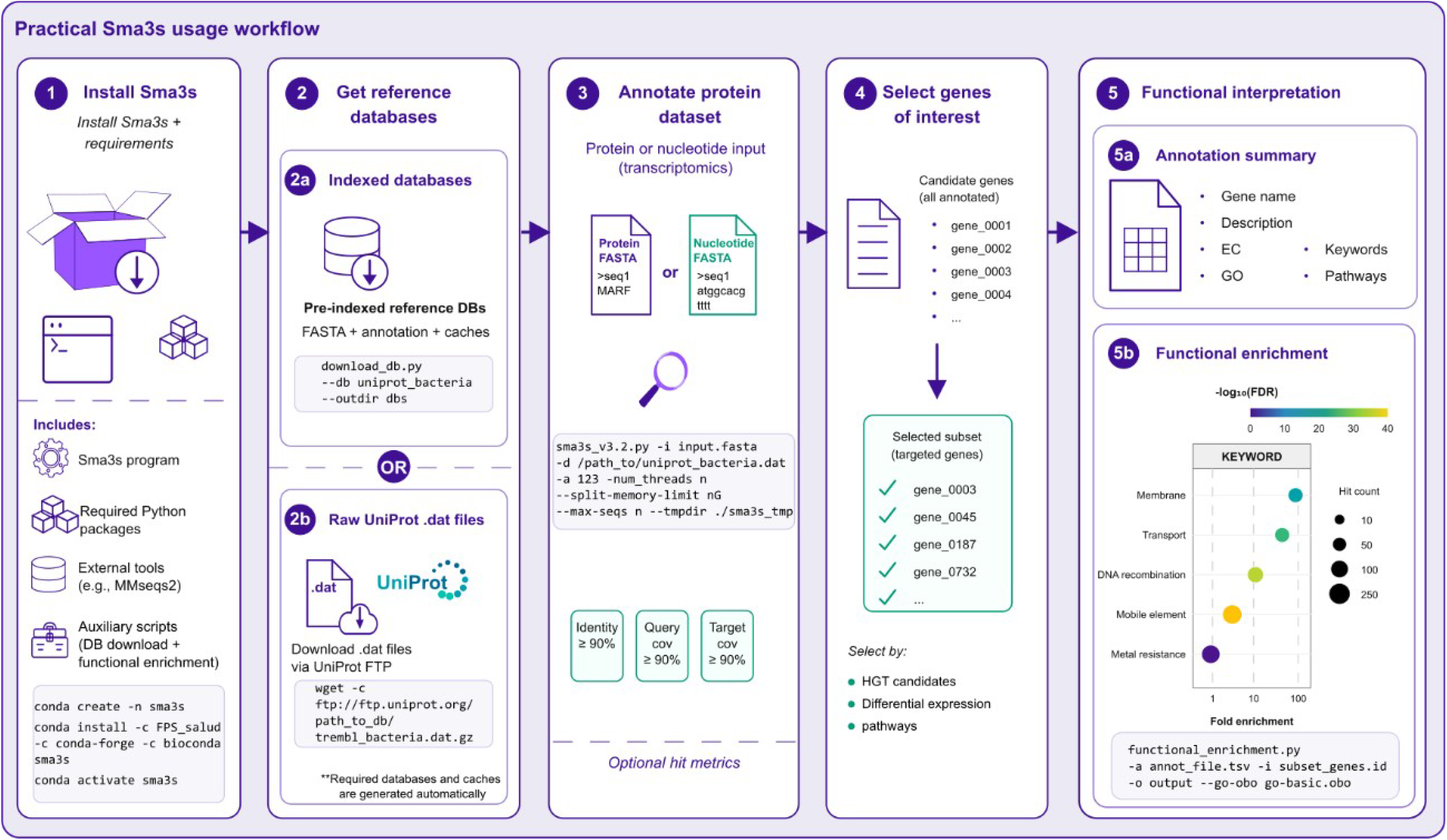
Sma3s v3 Workflow. This diagram provides a synopsis of the primary steps involved in the installation and utilization of Sma3s v3 in the context of reproducible functional analyses. The initial step entails the installation of the Conda package, which encompasses the primary program, ancillary dependencies such as MMseqs2, and auxiliary scripts. The subsequent step entails the preparation of the reference databases, which can be utilized as pre-indexed databases or generated from files with UniProt entries. After that, Sma3s generates the FASTA files, annotations, and caches necessary to expedite subsequent operations. The third step involves the annotation of input sequences, whether they be protein or nucleotide, using configurable thresholds for identity, coverage of the query sequence, and coverage of the target sequence. The fourth step delineates the selection of genes or clusters of interest from the annotation table. These may include, but are not limited to, candidate genes for horizontal gene transfer, differentially expressed genes, or genes associated with specific pathways. The fifth step encompasses the functional interpretation, which includes the generation of annotation summaries such as gene name, description, EC, GO, keywords, and pathways. It also involves functional enrichment analysis based on GO terms or UniProt keywords.

Sma3s v3 offers a comprehensive workflow that integrates the establishment of reference databases with the functional interpretation of gene sets, presented in a highly simplified and user-friendly manner. The analysis starts with pre-indexed databases or files with UniProt entries, from which the program generates the sequences, annotations, and caches necessary to reuse the same reference in subsequent analyses. Then, the input sequences are annotated using configurable identity and coverage thresholds, and the metrics of the hit responsible for each annotation can be retained. The primary output can be integrated with lists of genes of interest that are defined externally, thereby enabling the application of Sma3s in diverse biological contexts. These include the characterization of candidate genes for horizontal gene transfer, the identification of differentially expressed genes, and the analysis of subsets associated with specific pathways. In the final step of the process, the auxiliary scripts included in the distribution enable users to generate functional summaries and enrichment analyses based on GO terms or UniProt keywords (Figure 2).

### Sma3s v3 exhibits broad and semantically robust functional annotation compared to reference annotators

To assess the relative performance of this new Sma3s version against other widely utilized functional annotators within the scientific community, a *Vibrio cholerae* pangenome was constructed using genomes obtained from NCBI Genome. Prior to the inference of the pangenome, the initial dataset (comprising 16,618 genomes) underwent a quality filtration process to eliminate duplicate genomes, divergent assemblies, or genomes that were potentially misclassified. This procedure yielded a refined set of *V. cholerae* genomes (11,295 genomes) and reduced the inclusion of artifacts resulting from fragmented or taxonomically inconsistent assemblies by approximately 32% of the initial genomes. Then, this pangenome was functionally annotated using Sma3s, InterProScan^1^, and eggNOG-mapper^18^.

The comparative analysis of functional annotations unearthed discernible discrepancies among tools, contingent upon the filtration of genes bearing non-informative annotations, such as "hypothetical protein" or "unknown function". Sma3s v3 and InterProScan yielded highly comparable functional coverage, with Sma3s v3 annotating 30,662 genes, accounting for 60.8% of the pangenome, and InterProScan annotating 30,342 genes, equivalent to 60.2%. In contrast, eggNOG-mapper annotated 20,848 genes, representing 41.4% of the total analyzed, showing more limited functional coverage of the pangenome (Figure 3A). Subsequently, a functional overlap analysis was performed among the annotators to identify genes for which functional information was recovered. The analysis revealed that 18,176 genes exhibited concordant functional information derived from the three tools. It is noteworthy that both InterProScan and Sma3s had a substantial number of genes annotated exclusively by each annotator (4,153 and 5,747, respectively) and annotated jointly (6,229 genes). In contrast, the exclusive contribution of eggNOG-mapper was more modest, with a mere 378 genes annotated exclusively by this tool. Notably, more than 13,000 genes lacked valid functional evidence in any of the three annotators (Figure 3B).

**Figure 3.**
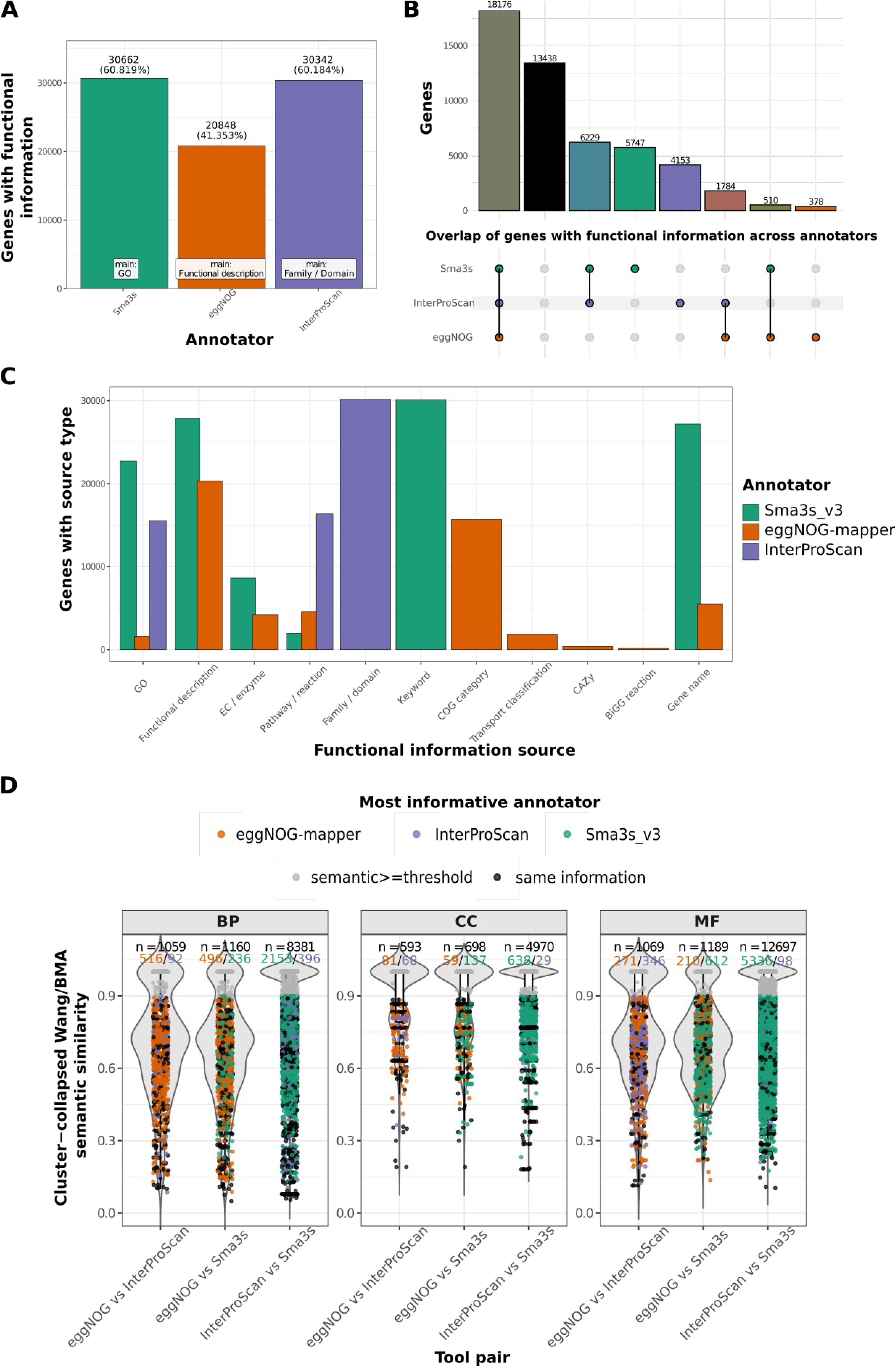
Functional and semantic comparison of annotators in the *Vibrio cholerae* pangenome. A) Number and percentage of pangenome clusters with at least one valid source of functional evidence from each tool after filtering out non-informative annotations. B) Overlap of clusters that have been functionally annotated by SMA3S v3, eggNOG-mapper, and InterProScan. The vertical bars in the plot represent the number of clusters in each intersection, while the horizontal bars indicate the total number of clusters annotated by each tool. C) Breakdown of the sources of functional evidence retrieved by each annotator. The categories were counted non-exclusively; therefore, a single cluster could contribute to multiple functional sources within the same tool. D) Comparison of semantic similarity among annotators based on GO terms collapsed into semantic clusters. Comparisons were performed in pairs and independently for Biological Process (BP), Cellular Component (CC), and Molecular Function (MF). The Wang/BMA similarity is a metric that quantifies the semantic agreement between the sets of GO terms assigned by two tools to the same cluster. Colored dots indicate the annotator that contributed the greatest number of effective semantic clusters for genes exhibiting a similarity below the exploratory threshold of 0.90; conversely, the gray dots indicate that the semantic difference between the two annotators is not significant.

The overlap among tools did not reveal the nature of the functional evidence recovered. Therefore, the types of evidence sources contributing to the annotations generated by each tool were examined (Figure 3C). The categorization of proteins was not performed in an exclusive manner; therefore, a given gene could contribute to multiple functional evidence sources within a particular tool. Sma3s v3 demonstrated a comprehensive annotation profile, predominantly comprising functional descriptions, UniProt keywords, gene names, GO terms, and EC numbers. InterProScan exhibited the hallmark characteristics of a signature-based annotation tool, manifesting a preponderance of family/domain assignments, accompanied by associated GO terms and functional pathways. Finally, eggNOG-mapper annotations were found to comprise several information sources, including functional descriptions, COG categories, GO terms, and other tool-specific sources (Figure 3C).

Given that GO terms constitute a common functional source for Sma3s v3, eggNOG-mapper, and InterProScan, they were used to assess semantic agreement among the tools beyond the simple presence or absence of functional annotation. However, the number of genes with GO terms varied among annotators, particularly in eggNOG-mapper. Therefore, comparisons were performed only on genes that had GO annotations in both tools for each pairwise comparison. This approach precluded the imposition of penalties on an annotator for the absence of GO terms, thereby enabling the evaluation of semantic similarity exclusively when both annotators provided comparable information.

To ascertain whether the observed discrepancies between the tools were attributable exclusively to the number of assigned GO terms or whether they reflected actual functional disparities, redundant GO terms were consolidated into semantic clusters, and the inter-annotator similarity was evaluated based on these reduced sets. The comparison was performed independently for Biological Process, Cellular Component, and Molecular Function. After semantic clustering, a large proportion of genes exhibited elevated Wang/BMA similarities across diverse tools. The Wang metric quantifies the semantic similarity between pairs of GO terms^31^, while BMA (Best Match Average) encapsulates the similarity between sets of GO terms.

These findings suggest that numerous GO annotations exhibited functional similarity despite being derived from different annotators. However, subsets of genes exhibiting similarity below the exploratory threshold of 0.90 were also identified. In these cases, the number of effective semantic clusters contributed by each annotator was compared to identifying which tool provided the greatest amount of non-redundant GO information (Figure 3D).

Among genes exhibiting a Wang/BMA similarity score below 0.90, the contribution of effective semantic information exhibited variability depending on the ontology and the comparison between tools. In the ontology of biological process, eggNOG-mapper yielded more effective semantic clusters than InterProScan for 516 genes and Sma3s for 496 genes, while Sma3s clearly outperformed InterProScan for 2,153 genes. In the context of molecular function, Sma3s demonstrated the most significant contribution in comparison to InterProScan, exhibiting enhanced clusters for 5,336 genes. Additionally, it surpassed eggNOG-mapper for 612 genes, underscoring its efficacy in gene annotation. In the Cellular Component category, the disparities were less pronounced, although Sma3s maintained a higher effective semantic contribution in comparisons against both eggNOG-mapper and InterProScan. The results of this study suggest that semantic discrepancies are not uniformly distributed across ontologies. Furthermore, Sma3s has been found to provide a greater amount of non-redundant GO information in comparison to InterProScan, particularly within the categories of Molecular Function and Biological Process.

### Application of Sma3s v3 to large-scale pangenome annotation and screening for horizontal gene transfer (HGT) signals

Once the effectiveness of Sma3s had been demonstrated, the resulting functional annotation was analyzed. The pangenome of *V. cholerae* comprised 50,415 gene clusters, of which 2,837 were classified as core, 35,450 as accessory genome, and 12,128 as unique. Consequently, the majority of the pangenome was concentrated in the accessory compartment, while approximately one-quarter of the clusters corresponded to unique or low-prevalence genes.

The integration of annotations generated by Sma3s exhibited a discernible dependence on the pangenome category under analysis. Within the core group, 2,755 of 2,837 clusters were annotated, constituting 97.1% of the group. Conversely, the proportion of annotated clusters in the accessory genome was lower, with 20,719 out of 35,450 clusters annotated (58.4%), and in the unique fraction, with 7,188 out of 12,128 clusters annotated (59.2%) (Figure 4A). These results suggest that Sma3s recovered functional annotations almost completely for conserved genes, while a substantial proportion of variable or strain-specific genes lacked detectable functional annotation compared to the reference used.

**Figure 4.**
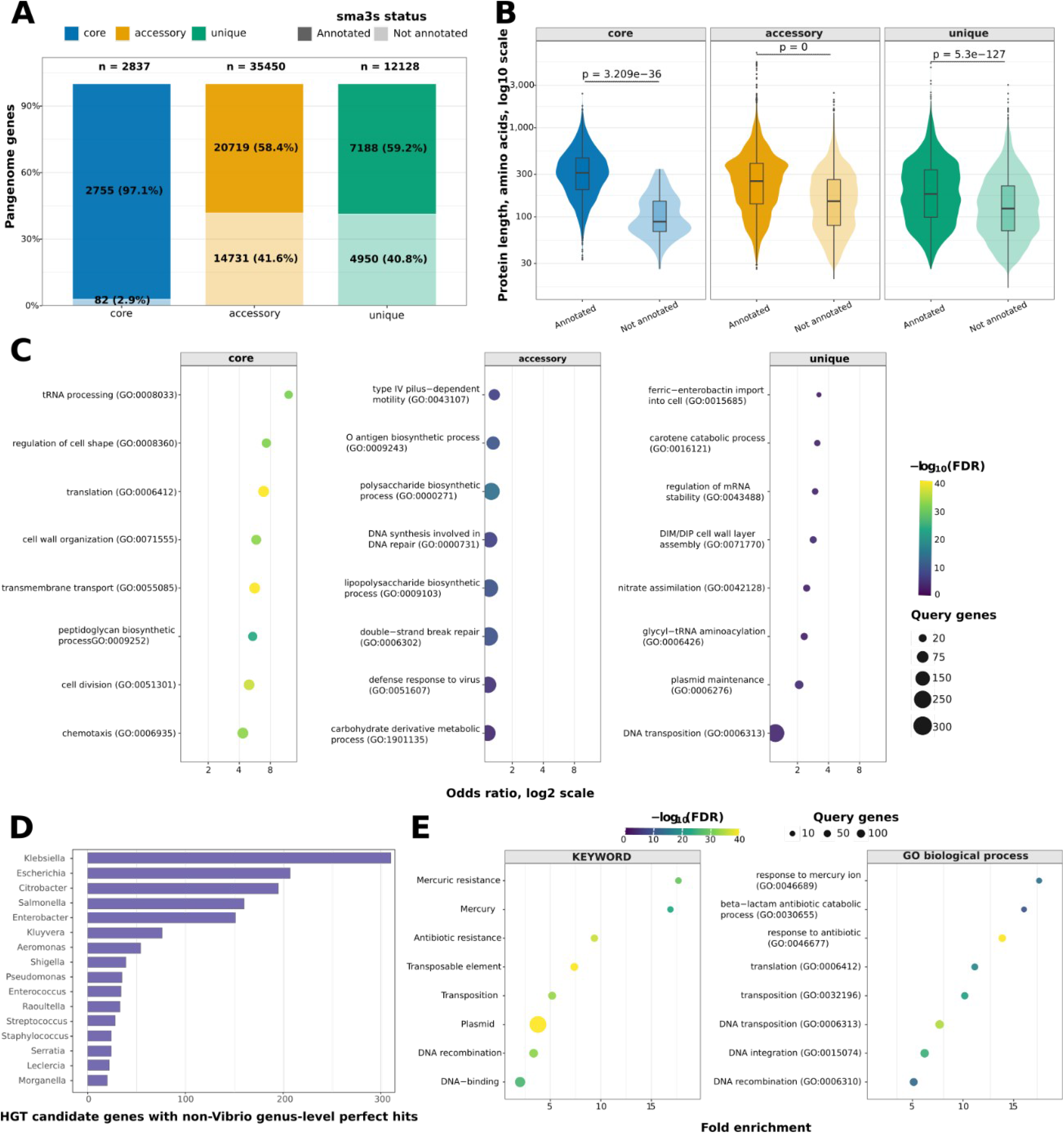
Sma3s v3 annotation and functional enrichment of the *V. cholerae* pangenome. A) Proportion of gene clusters annotated and unannotated by Sma3s v3 across the core, accessory, and unique pangenome categories. Bars show the percentage of annotated and unannotated clusters within each category, whereas the labels inside the bars indicate the absolute number and corresponding percentage of clusters. The total number of clusters in each category is shown above the bars. B) Distribution of reference protein sequence lengths for annotated and unannotated clusters in each pangenome category. Protein length is shown on a logarithmic scale. Differences between annotated and unannotated proteins within each category were assessed using the Wilcoxon rank-sum test. C) Enrichment of Gene Ontology Biological Process terms in the core, accessory, and unique categories of the *V. cholerae* pangenome. For each category, enriched terms were identified relative to the remainder of the pangenome using a one-sided Fisher’s exact test followed by false discovery rate (FDR) correction. The x-axis shows the log2 odds ratio, point color indicates statistical significance as −log10(FDR), and point size represents the number of clusters associated with each term. D) Non-*Vibrio* genera most frequently associated with candidate horizontally transferred genes, defined as pangenome genes with protein matches exhibiting 100% sequence identity and full-length coverage against sequences from genera outside *Vibrio*. Bars indicate the number of candidate genes with matches assigned at the genus level. E) Functional enrichment of candidate horizontally transferred genes based on UniProt Keywords and GO Biological Process terms. The x-axis shows fold enrichment relative to the complete pangenome background. Point size indicates the number of candidate genes associated with each term, and point color represents adjusted statistical significance as −log10(FDR).

To investigate the association of the absence of annotation with structural characteristics of the proteins, the lengths of annotated and unannotated sequences were compared within each pangenome group. Annotated proteins were consistently longer than unannotated proteins across all three categories, with significant differences observed in the core, accessory, and unique groups (Wilcoxon rank-sum test; core, p = 3.209 × 10^-36^; accessory, p=0; unique, p = 5.3 × 10^-127^; Figure 4B). This tendency manifested to a particularly marked degree in the core group, wherein the limited number of unannotated proteins exhibited a marked reduction in size.

Functional enrichment analysis yielded discernable profiles among the various groups of the *Vibrio cholerae* pangenome (Figure 4C). With respect to GO biological processes, core genes were enriched in conserved functions, including tRNA processing, translation, transmembrane transport, cell division, cell shape regulation, and chemotaxis. The accessory gene group was enriched in processes related to surface structures and genomic variability, such as the biosynthesis of polysaccharides, O-antigen, and lipopolysaccharides; type IV pilus-dependent motility; DNA repair; and the viral response. The unique group showed enrichment in low-prevalence processes associated with genetic mobility and highly specific functions, including DNA transposition, plasmid maintenance, ferric-enterobactin import, and nitrate assimilation. This pattern was consistent with complementary analyses based on UniProt keywords and GO terms for molecular function, in which the “core” fraction was associated with conserved cellular functions, the “accessory” fraction with glycosyltransferases, nucleases, DNA integration, and antiviral defense, and the “unique” fraction with plasmids, motility proteins, conjugation, and transposase activity (**Supplementary Figure XX**).

To assess the utility of Sma3s v3 for identifying signals consistent with horizontal gene transfer in bacterial pangenomes, *V. cholerae* gene clusters were systematically screened for exact amino acid matches to proteins from non-*Vibrio* genera, requiring 100% sequence identity and full-length coverage. Under this stringent criterion, 1,838 candidate genes were identified, all belonging to the unique fraction of the pangenome. These candidates represented 15.2% of the unique clusters and 3.6% of the complete pangenome, indicating that perfect matches to proteins from non-*Vibrio* genera were restricted to low-prevalence clusters detected in a single strain.

The non-*Vibrio* genera most frequently associated with these candidates were predominantly members of the Enterobacteriaceae family, particularly *Klebsiella*, *Escherichia*, *Citrobacter*, *Salmonella*, and *Enterobacter*. Outside this family, frequent matches were also observed with genera such as *Aeromonas* and *Pseudomonas* (Figure 4D). These genera should be interpreted as taxa containing proteins that perfectly matched the candidate genes, rather than necessarily representing their direct donors, because this screening criterion cannot determine the direction of transfer or distinguish recent horizontal gene transfer from alternative explanations for sequence sharing.

Functional enrichment analysis nevertheless revealed a profile consistent with genetic mobility and resistance to antimicrobial compounds and metals. Significantly enriched UniProt Keywords included terms related to plasmids, transposition, antibiotic resistance, and mercury resistance. Consistently, enriched Gene Ontology Biological Process terms included β-lactam antibiotic catabolism, response to mercury ions, and DNA transposition (Figure 4E). Together, these findings suggest that exact-match screening against proteins from non-*Vibrio* genera identifies a functionally distinct subset of the unique pangenome enriched in functions related to genetic mobility, antimicrobial resistance, and metal tolerance.

### Sma3s v3 increases coverage and functional diversity in a large-scale metagenomic catalog

To evaluate SMA3S in an applied metagenomic scenario, the MGYA00131824 analysis from MGnify was selected. This analysis is derived from the MGYS00001842 study, which focused on the detection and strain-level identification of Shiga toxin-producing *Escherichia coli* in contaminated spinach samples^36^. This dataset offers an ideal use case, as it encompasses a complex microbial community, with an anticipated signal from *E. coli* and the Enterobacteriaceae family, in addition to proteins that have been previously annotated through the utilization of an earlier iteration of InterProScan.

The protein catalog that was downloaded from MGnify contained 843,935 sequences, of which 535,830 corresponded to proteins that had been previously annotated and 308,105 to proteins that had not been annotated in the original analysis. As these data had initially been processed using an earlier version of InterProScan, version of 2017 analysis, the entire catalog was subsequently re-annotated using the most recent version of InterProScan and Sma3s. This strategy enabled the evaluation of Sma3s’s capacity to process an entire metagenomic dataset and the comparison of its performance against domain/family-based annotations.

Sma3s functionally annotated 728,014 proteins, equivalent to 86.3% of the catalog, outperforming both InterProScan version of 2026 (616,895 proteins; 73.1%) and the original InterProScan version of 2017 annotation (494,489 proteins; 58.6%) (Figure 5A). Furthermore, Sma3s provided high coverage of specific functional layers, including keywords in 723,989 proteins (85.8%), gene names in 705,292 (83.6%), and GO terms in 697,696 (82.7%). In contrast, InterProScan 2026 was dominated by domain/family sources such as Gene3D, Pfam, and SUPERFAMILY, while the 2017 annotation showed more limited overall coverage, particularly for Gene3D and Prositepatterns. These results indicate that Sma3s substantially increases the number of proteins with retrievable functional information in a large-scale metagenomic catalog.

**Figure 5.**
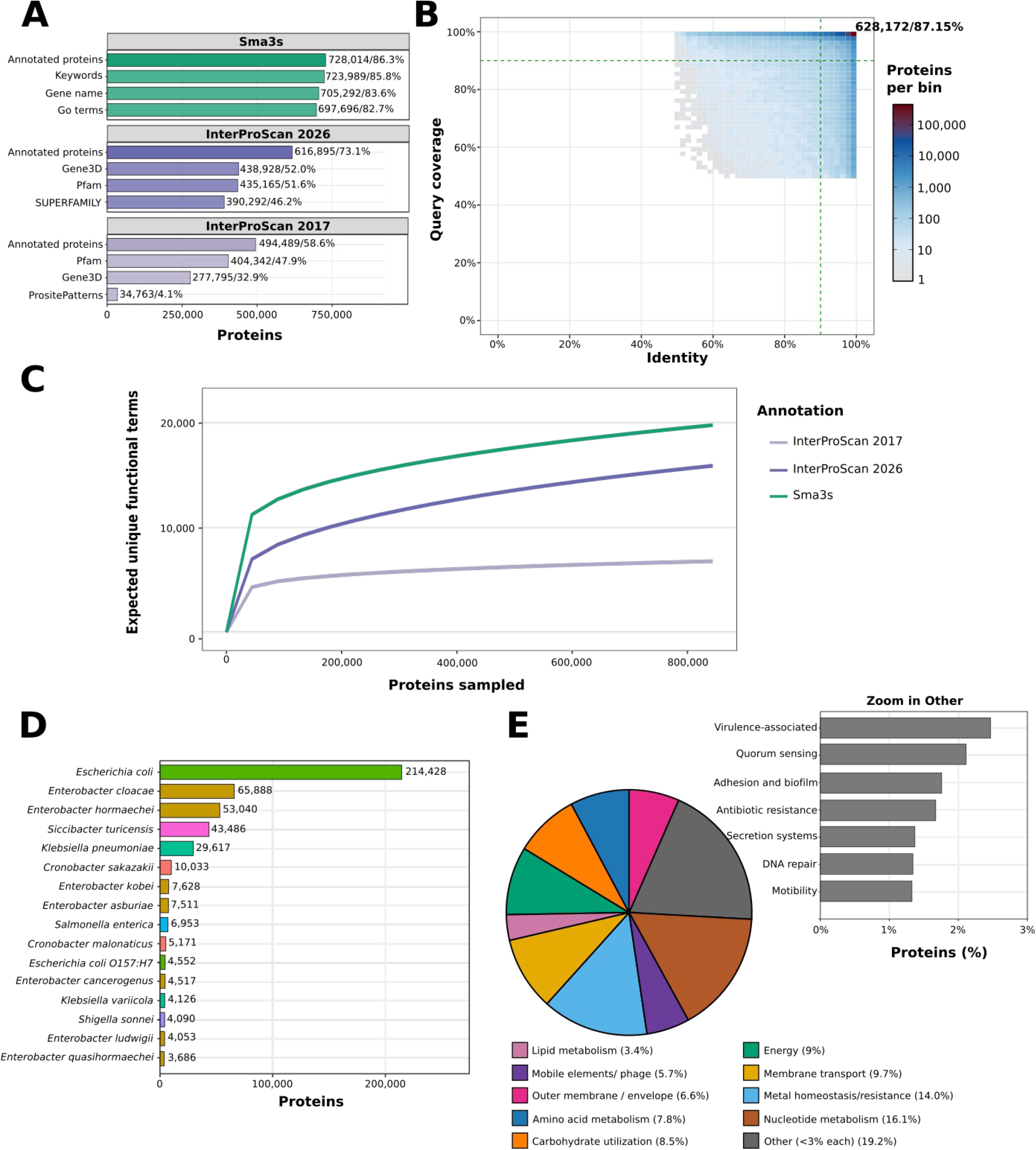
Sma3s v3 enables scalable functional annotation of a food-contamination metagenomic catalog. A) Number of proteins annotated by Sma3s, InterProScan 2026, and InterProScan 2017, together with the main information sources or annotation layers recovered by each method. B) Joint distribution of amino acid sequence identity and query coverage for the best Sma3s hits against the reference database. Only hits with ≥50% identity and ≥50% coverage are shown; dashed lines indicate the high-confidence threshold, defined as ≥90% identity and ≥90% coverage. C) Analytical rarefaction curves showing the expected number of unique functional terms recovered as the number of sampled proteins increased. D) Species associated with the largest numbers of high-confidence Sma3s hits. Counts represent proteins whose best hit was assigned to each species. E) Distribution of functional categories inferred from Sma3s annotations. Categories account for <3% of the annotated proteins were grouped as Other. This category was further subdivided to show less frequent functions associated with virulence, signaling, biofilm formation, antibiotic resistance, secretion systems, DNA repair, motility, and defense systems.

The robustness of the annotations generated by Sma3s was assessed from the joint distribution of amino acid sequence identity and query coverage for the best hits against the reference database. Most hits were concentrated in the high-identity, high-coverage region (Figure 5B), with 628,172 sequences falling within the quadrant defined by ≥90% identity and ≥90% coverage, corresponding to 87.15% of the displayed hits. The remaining annotated sequences were distributed across regions of lower identity and/or coverage, showing that Sma3s also recovered functional annotations from more divergent or partial matches.

The functional diversity recovered by each annotator was then compared using analytical rarefaction curves of unique functional terms. This analysis enabled the estimation of the expected increase in the functional repertoire as more proteins from the metagenomic catalog were progressively incorporated. Sma3s demonstrated a more substantial functional recovery across the entire range that was analyzed, reaching approximately 20,000 unique functional terms when the complete catalog was taken into consideration. InterProScan 2026 showed intermediate recovery, while InterProScan 2017 exhibited the lowest functional diversity (Figure 5C).

The taxonomic signal associated with the proteins annotated by Sma3s was explored based on the best high-confidence hits defined by ≥90% identity and ≥90% coverage. The species with the highest number of associated proteins was *Escherichia coli* (214,428 proteins), followed by *Enterobacter cloacae* (65,888), *Enterobacter hormaechei* (53,040), *Siccibacter turicensis* (43,486), and *Klebsiella pneumoniae* (29,617) (Figure 5D). These values signify the number of proteins for which the most reliable match was associated with each species, not a direct estimation of taxonomic abundance.

The final analysis of the functional categorization of the proteins annotated by Sma3s revealed a profile consistent with a bacterial metagenome associated with food contamination (Figure 5E). The categories with the highest representation included nucleotide metabolism, homeostasis/metal resistance, membrane transport, energy metabolism, carbohydrate utilization, amino acid metabolism, and outer membrane/envelope components. A subsequent breakdown of the “Other” category (representing less than 3% of the total) yielded several noteworthy functional categories, including those associated with virulence, quorum sensing, adhesion/biofilm, antibiotic resistance, secretion systems, DNA repair, and motility. Taken together, these results show that Sma3s can be applied on a scalable basis to a metagenomic catalog of more than 840,000 proteins. This application increases both the number of proteins with functional annotations and the diversity of functional terms retrieved. This increase is compared to the original 2017 annotation and the updated InterProScan.

## Discussion

Functional annotation remains a primary challenge in the interpretation of genomic and metagenomic data. Current catalogs may contain hundreds of thousands or millions of proteins, so annotation methods must combine sensitivity, scalability, and traceability. In this paper, we present a new version of Sma3s that retains the modular strategy of its previous versions^3,4^, but incorporates an architecture adapted for processing large sets of sequences. The incorporation of MMseqs2 and the parallelization of the workflow reduce the reliance on strictly sequential steps and allow for the efficiency of large-scale protein similarity searches to be leveraged^24^. These enhancements broaden the scope of Sma3s, expanding from individual proteomes to pangenomes and large-scale metagenomic catalogs.

The value of this new version, Sma3s v3, lies not only in increasing the number of annotated proteins, but also in doing so while maintaining explicit information about the evidence supporting each assignment. Consequently, in the tool’s stress tests employing pan-genomic and metagenomic data, the outcomes were substantiated by significant alignments with high identity and coverage values. Assignments derived from more distant homologies persisted in their identifiable state through the utilization of similarity metrics. This traceability is of particular importance because the reliability of functional transfer depends not only on the existence of a significant hit, but also on the identity and coverage of the alignment, membership in a functionally coherent family or subfamily, and the level of specificity of the transferred annotation^37^. Therefore, this feature of Sma3s v3 is essential because current databases include both manually reviewed records and annotations generated by computational methods. In contrast to the assumption that all matches provide an equivalent level of evidence, Sma3s provides the necessary information to evaluate, filter, and audit each functional transfer, if deemed necessary^1,25,38^.

In this context, Sma3s v3 can be utilized conceptually at two complementary levels of confidence. A conservative mode, predicated on high identities and coverage, would be congruent with applications that prioritize minimizing uncertain assignments, such as those employed in the screening of determinants associated with resistance, virulence, or genetic mobility. Conversely, an exploratory mode could retain assignments based on more distant homologies to investigate poorly characterized proteins and expand the recovered functional space, provided that these are considered hypotheses subject to further validation^39^. This flexibility precludes the implementation of a uniform threshold across disparate biological targets, thereby enabling the modulation of the equilibrium between coverage and specificity. The relationship between these two factors is contingent upon the protein family and the level of functional intricacy sought to be captured. A semantic comparison with other annotators lends support to the utility of this exploratory approach, as Sma3s assignments maintained overall functional consistency even when they provided information not recovered by InterProScan or eggNOG-mapper. These results carry significant weight due to the inherent limitation of a binary annotation approach, which is inadequate for evaluating annotators. This limitation stems from the potential for disparate tools to assign analogous numbers of GO terms yet exhibit marked disparities in terms of redundancy, specificity, and biological consistency. Consequently, Sma3s enhances the quantity of retrieved information, which occupies a functional space consistent with that described by independent methods. However, it is imperative to acknowledge that this agreement should be regarded as an indication of computational robustness rather than a validation of the accuracy for each protein. CAFA exercises have demonstrated that the automated function prediction domain remains to be significantly enhanced, even for the most efficacious methods^40^.

Our results also demonstrated that the functional diversity recovered by Sma3s is complementary to that obtained using InterProScan and eggNOG-mapper. InterProScan employs models from multiple databases to identify domains, families, and conserved sites^1^. In contrast, eggNOG-mapper utilizes orthology assignments to transfer functional information^18^. Sma3s extracts an integrated repertoire of gene names, GO terms, EC numbers, keywords, and pathways based on reference proteins. The enhanced diversity observed when combining Sma3s and InterProScan suggests that these approaches delineate partially distinct components of protein function. Similar evidence-integration strategies have been implemented in tools such as Mantis, which derives consensus annotations from concordant information across multiple reference sources^41^. Consequently, Sma3s v3 can function as both a primary annotator and a complementary functional layer within multi-annotator workflows, while providing a basis for the future development of a consensus score integrating different types of evidence. Assignments independently supported by sequence similarity, orthology and conserved-domain detection could receive higher confidence than those based on a single source, whereas discrepancies between tools could identify proteins requiring individual review. Sma3s v3 is well suited to this integration because it preserves similarity metrics and functional categories that can be traced back to the original evidence.

Metagenomic analysis highlighted an additional application of Sma3s v3: expanding the functional interpretation of previously processed catalogues. The dataset analyzed in this study was derived from commercial spinach experimentally inoculated with the Shiga toxin-producing *E. coli* strain EC1276 (Sakai). Therefore, the prominent *E. coli* signal is consistent with the experimental design and does not necessarily reflect its natural abundance in the spinach microbiota^36^. Likewise, the taxonomic distribution of the best protein hits reflects similarities to reference proteins and should not be equated with an abundance profile derived directly from sequencing reads using specialized tools such as MetaPhlAn 4^42^. Furthermore, the recovery of approximately 20,000 unique functional terms, together with the position of the Sma3s curve above those of InterProScan 2017 and InterProScan 2026, support that Sma3s substantially expands the functional repertoire recovered. Taken together, these results show that Sma3s can transform large metagenomic catalogues into interpretable information and facilitate the prioritization of functions related to bacterial metabolism, the cell envelope, genetic mobility, adaptation, biofilm formation and resistance.

Beyond this biological interpretation, comparison of the original annotations generated with InterProScan 2017 with those obtained using current versions shows that functional annotation depends on when it is performed and should therefore be regarded as a dynamic process. This temporal dimension has been recognized by resources such as RefSeq through the concept of “annotation age*”* and the implementation of continuous reannotation strategies designed to incorporate new evidence and methodological improvements^43,44^. Extending this concept, a protein catalogue has not only a sequencing date but also a “functional annotation age”, determined by the versions of the tools and databases used and, consequently, by the state of knowledge they captured at the time of analysis. The potential efficacy of Sma3s v3 in mitigating this issue lies in its capacity to directly re-annotate public protein catalogs. Resources such as MGnify^45^ consolidate vast quantities of analyses derived from particular iterations of their workflows. In instances where predicted proteins persist, their associated functional data can be revised without the necessity of reiterating quality control, assembly, and gene prediction processes. The scalability of Sma3s enables the direct reanalysis of these proteins against current databases, thereby reducing the cost of updating historical resources. The application could also be extended to longitudinal annotation monitoring. The periodic reannotation of the same catalog would make it possible to quantify which proteins transition from uncharacterized to annotated and which new functions are incorporated as knowledge advances. So, Sma3s could be utilized not only to generate a one-time annotation but also to assess the temporal progression of available functional information.

Analysis of the *V. cholerae* pangenome shows, however, that database growth and the combined use of multiple annotators do not fully resolve the lack of functional information, as approximately one-quarter of the pangenome, 13,000 of the 50,415 gene clusters, received no informative annotation from any of the tools used in this study. This fraction was concentrated primarily in the accessory and unique groups, which contain less conserved genes, genes restricted to particular strains, and genes that are poorly represented in reference databases. This finding is consistent with the open structure of the *V. cholerae* pangenome, which is characterized by extensive accessory gene diversity^46^. This feature, together with the shorter length of many of the corresponding proteins, reduces the sequence signal available for detecting homologs or complete domains. Microbiome studies have shown that small proteins are underrepresented by conventional prediction and annotation systems, despite including conserved and potentially functional families^47,48^. At the same time, rare pangenome’s group may contain fragments, gene prediction errors, or spurious sequences, particularly in pangenomes constructed from thousands of genomes^30^. The absence of an annotation should therefore be interpreted as insufficient evidence rather than necessarily indicating an absence of function. Collectively, these observations highlight that the main challenge is no longer limited to processing large numbers of sequences, but also involves functionally interpreting “rare” genes that fall outside current knowledge of common or essential genes. Recurrence could be used to prioritize this functionally uncharacterized fraction. An unannotated protein detected in multiple strains, conserved within a syntenic context, or independently recovered across different catalogs has stronger support as a genuine biological product than an isolated singleton sequence. Sma3s could facilitate this process by enabling the joint reanalysis of large protein collections and the grouping of proteins that remain functionally uncharacterized but display recurrent distributions. This approach would transform the unannotated fraction into a prioritized set of candidates for subsequent characterization. The application of Sma3s v3 to horizontal gene transfer screening illustrates its potential for generating evolutionary hypotheses. The exclusive occurrence within the unique fraction of all 1,838 genes showing perfect matches and full-length coverage against proteins from non-*Vibrio* genera is consistent with recent or sporadic acquisitions that remain narrowly distributed within the species. Identical sequences shared by distantly related bacteria have been used as evidence of recent genetic exchange because complete identity becomes progressively less likely as the time since sequence divergence increases^49^. Some of the observed associations are also ecologically plausible. In particular, members of *Aeromonas* and *Vibrio*, including *V. cholerae*, can coexist in aquatic environments and colonize shared chitin-associated hosts or substrates, such as planktonic copepods and chironomid egg masses^50,51^. This ecological overlap could provide opportunities for genetic exchange. However, perfect matches do not allow the direction of transfer to be inferred, a direct donor to be identified, or alternative explanations, such as extreme sequence conservation, contamination, or taxonomic misassignment, to be completely excluded. These genes should therefore be regarded as candidates consistent with horizontal gene transfer rather than as evidence of confirmed transfer events.

The functional enrichment of these candidates provides an additional line of evidence supporting their association with genetic mobility. Functions related to plasmids, transposition, recombination, and antimicrobial resistance are consistent with the established contribution of mobile genetic elements to the evolution of *V. cholerae*. SXT/R391 elements have been shown to comprise a well-defined family of integrative and conjugative elements that can mobilize resistance determinants in this species^52,53^. Concomitantly, the co-occurrence of antibiotic- and metal-resistance genes may promote co-selection^54^. Collectively, the results indicate Sma3s v3 possesses the capability to integrate functional annotation with the taxonomic distribution of protein matches, thereby facilitating the narrowing of a complex pangenome to a manageable set of prioritized genes and functions. In the future, the incorporation of genomic context, the identification of nearby mobile genetic elements, and the reconstruction of gene phylogenies will enable a more robust assessment of the origins and evolutionary histories of these candidates.

The main limitations of Sma3s v3 are inherent to the processes of homology-based annotation and functional transfer. The comprehensiveness and precision of annotations are contingent upon the taxonomic and functional representation of the reference databases, as well as the heterogeneity between reviewed and computational annotations. Despite being common to all homology-based methods, these limitations do not invalidate the resulting assignments. Rather, they require interpretation in light of sequence identity, alignment coverage, and evidence provenance, in addition to the application of filters tailored to the objective of each analysis. The potential for future investigation of proteins that remain unannotated lies in the implementation of structure-based approaches. Tools such as Foldseek facilitate the expeditious comparison of predicted structures against voluminous reference collections^55^. Global metagenomic analyses have demonstrated that structural similarity can reveal potential functional relationships that are undetected by conventional sequence searches^56^. This strategy could offer insights into the evolution of divergent proteins within a flexible genome, contingent upon the reliability of the corresponding structural models.

Protein language models offer an alternative approach for expanding functional coverage. Tools such as DeepGO-SE and ProtNote leverage protein embeddings to predict GO terms, enzyme activities, or protein–function associations^23,57^, whereas PLMSearch uses these representations to detect remote homology^58^. However, these methods are often restricted to specific functional spaces, and their performance cannot be directly extrapolated to the comprehensive annotation of bacterial genomes. Many have been trained on datasets that inadequately represent microbial diversity and have undergone limited evaluation on complete genomes and complex metagenomes^59^. In the context of analyzing microorganisms, methods that are specifically tailored to this domain, such as DeepGOMeta^59^, or methods that are focused on the organization of bacterial genomes, such as SAFPred, which combines protein embeddings with conserved synteny^60^, are more appropriate.

The distribution of Sma3s v3 through the Anaconda ecosystem, in conjunction with the automation of database downloads and cache creation, enables its incorporation into reproducible protocols. This facilitates the tracking of versions of the dependencies and resources utilized^26^. In summary, the Sma3s has been upgraded to transform the original strategy into a scalable, flexible, and updatable system for the annotation and reannotation of large protein catalogs. Its primary contribution lies not only in expanding functional coverage but also in generating evaluable assignments that can be combined with evidence from domains, orthology, structure, or language models. These features position Sma3s v3 as a useful component for standardized annotation protocols for proteomes, pangenomes, and metagenomes, and for maintaining these resources in alignment with the evolution of biological knowledge.

